# Formation of multinucleated variant endothelial cells with altered mitochondrial function in cultured coronary endothelium under simulated diabetes

**DOI:** 10.1101/622407

**Authors:** Hilda Carolina Delgado De la Herrán, Luis Donis-Maturano, Carolina Álvarez-Delgado, Francisco Villarreal, Aldo Moreno-Ulloa

## Abstract

Coronary endothelial dysfunction is associated with atherosclerosis and myocardial infarction in subjects with type 2 diabetes mellitus (T2DM). Vascular endothelial cells are referred to as small and polygonal mononuclear cells. However, multi-nucleated and large endothelial cells (named as multinucleated variant endothelial cells [MVECs]) have been reported in the aorta, wherein their abundance correlates with atherosclerosis severity. The role of MVECs in coronary endothelium remains obscure. We hypothesized that simulated diabetic conditions increase the number of MVECs and affect their mitochondrial structure/function in cultured coronary endothelium. The *in vitro* model of diabetes consisted in the treatment of bovine coronary artery endothelial cells (BCAECs) with high-insulin (100 nmol/L, HI) for three days followed by high-glucose (20 mmol/L, HG) and HI for nine additional days. Simulated diabetic conditions increased the abundance of MVECs compared to normal glucose (NG, 5.5 mM). MVECs had a higher nucleic acid content (7.2-Fold), cell diameter (2.2-Fold), and cell area (11.4-Fold) than mononuclear cells. Immunodetection of von-Willebrand factor (endothelial cell marker) in MVECs was positive. The mitochondrial mass was reduced, and mitochondrial membrane potential increased in mononuclear cells cultured in HG+HI compared to mononuclear cells grown in NG. However, the opposite mitochondrial findings were noted in MVECs compared to mononuclear cells. Mass spectrometry-based quantitative proteomic and gene ontology analysis suggested augmented mitochondrial autophagy, apoptosis, and inflammation suppression in cells cultured under HG+HI compared to NG conditions. These findings show that simulated diabetes increases the abundance of MVECs, and that mitochondrial structure and function are differentially affected between MVECs and mononuclear cells.

## 1. Introduction

The vascular endothelium is an organ that contributes to the proper functioning of the circulatory system [1]. In the heart, damage to coronary artery endothelial cells leads to coronary endothelial dysfunction, which is associated with the development of cardiac pathologies [2]. Particularly, subjects with type 2 diabetes mellitus are at an increased risk of myocardial infarction [3], and coronary endothelial dysfunction has been implicated in the process [4]. Coronary endothelial dysfunction is an independent predictor of future acute cardiovascular events in subjects with and without atherosclerosis [2]. Noteworthy, the majority of studies assessing coronary endothelial function have focused on endothelium-dependent vasodilation [5] and intracellular cell signaling pathways related to nitric oxide (i.e., *in vitro*) [6], but less is known beyond this established research panorama.

Endothelial cells are commonly referred to as small polygonal and mononuclear cells displaying a cobblestone-like monolayer when grown under static *in vitro* conditions [7, 8]. Although, the endothelial shape is influenced by various physiological and physical factors (e.g., shear stress, matrix support, etc.) [9-11] they remain mono-nucleated during the resting cellular stage as observed under *in vitro* conditions. It is recognized that there exists heterogeneity in the endothelium throughout the circulatory system [12]. The discovery of heterogeneity (e.g., function and organization) in this organ opened a conundrum about the differential functions of endothelial cells among organs and tissues and their role in specific diseases.

However, studies have reported the appearance of a distinct endothelial phenotype in the aorta of subjects with atherosclerosis wherein its presence correlates with the severity of the disease [13]. Also, this type of cells presented with altered functions including augmented LDL uptake [14], thus hinting at a plausible role for this phenotype in the development of cardiovascular disease (CVD). These cells are multinucleated (>2 nuclei) and larger than regular endothelial cells and, when first reported, were referred to as multinucleated variant endothelial cells (MVECs) [13]. The presence and function of MVEC in other vascular beds remain obscure principally in those organs intrinsically involved in the development of CVD.

We hypothesized that high-glucose (HG) and high-insulin (HI) (simulated diabetes) lead to the formation of MVECs in cultured coronary artery endothelial cells and that MVECs present with altered mitochondrial structure or function. Here, we evaluated the appearance and abundance of MVECs after prolonged exposure (9 days) to HG and HI and characterized their morphology, as well as their mitochondrial mass and function. In addition, we performed a quantitative proteomic differential analysis among endothelial cells cultured under normal-glucose (NG) and HG+HI. Overload of glucose and insulin led to an increased abundance of MVECs with a higher nucleic acid content, cell diameter, and cell area than mononuclear cells along with an augmented mitochondrial mass. Proteins involved in mitochondrial autophagy, apoptosis, and inflammation-suppression were upregulated in cells cultured with HG+HI compared to those in NG. Thus, our study provides evidence for the first time that simulated diabetes leads to the formation of MVECs with altered mitochondrial function in cultured coronary artery endothelium, which may contribute to vascular pathology as seen with diabetes.

## 2. Methods

### 2.1 Chemical and reagents

Recombinant human insulin was purchased from Sigma Aldrich (St. Louis, MO, USA). Antibiotic-antimitotic solution, trypsin-EDTA solution 0.25%, Hank’s Balanced Salt Solution (HBSS) without phenol red, Dulbecco’s Modified Eagle’s Media (DMEM) with glutamine, Fetal Bovine Serum (FBS), MitoTracker^®^ Green FM, MitoTracker^®^ Red FM, Hoechst 33258, Pentahydrate (bis-Benzimide)-FluoroPure™, methanol-free formaldehyde (16% solution), Pierce™ Quantitative Colorimetric Peptide Assay kit, and Pierce™ Mass Spec Sample Prep Kit for Cultured Cells were obtained from Thermo Fisher Scientific (Waltham, MA, USA). Acetonitrile, Optima™ LC-MS Grade, and water Optima™ LC-MS Grade were obtained from Fisher Scientific (Hampton, NH, USA). Rabbit anti-Von Willebrand factor (vWf) antibody and goat anti-rabbit IgG conjugated to Alexa Fluor 488 were from Abcam (Cambridge, MA, USA).

### 2.2 Cell culture

Bovine coronary artery endothelial cells (BCAECs) were purchased from Cell applications, Inc. (San Diego, CA, USA). For proliferative conditions, cells were grown with DMEM (5.5 mmol/L glucose, supplemented with 10% FBS and 1% antibiotic-antimitotic solution) at 37 °C in an incubator with a humidified atmosphere of 5 % CO_2_. When cells reached 70-80% confluence were split according to the manufacturer’s instructions. For simulated diabetic conditions, DMEM with 1% FBS was used to maintain cells under a quiescent mode. The scheme followed to simulate diabetes *in vitro* consisted of the following steps: 1) to induce insulin resistance, cells were treated for 3 days with 100 nmol/L insulin in normal glucose (NG, 5.5 mmol/L) DMEM [15]; then, 2) cells were maintained for 9 days with 100 nmol/L insulin in high glucose (HG, 20 mmol/L). This sequential scheme was followed to mimic the pathophysiological conditions that occur in T2DM, wherein hyperinsulinemia precedes hyperglycemia [16].

### 2.3 Abundance and morphological characterization of MVECs

Cells were seeded at 100 000 cells per well in 12-well plates (Corning^®^ CellBIND^®^) and treated as abovementioned. After HG and HI conditions, cells were washed with PBS to remove dead cells and debris. Two types of image analysis were performed using ImageJ software (version 2.0.0). MVECs were identified by the presence of more than two nuclei employing transmitted-light microscopy using 20x and 40x air objectives (EVOS™ XL Cell Imaging Station). Cell area and diameter were calculated by tracing lines (calibrated by image scale) around the cell perimeter and along the cell long axis, respectively. For visualization of nucleic acids, cells were stained with Hoechst 33258 (2 µg/ml in HBSS) for 30 minutes at room temperature (RT) and washed 3 x with PBS. Fluorescence images were taken using an EVOS^®^ FLoid^®^ Cell Imaging Station with a fixed 20x air objective. The quantification of the nucleic acid content was done by calculating the corrected total cell fluorescence (CTCF) tool from Image J software using the following equation: Integrated density-(Area of selected cell x mean fluorescence of background readings) [17]. At least three random fields per condition were chosen to capture images.

### 2.4 Immunofluorescence

Cells were seeded at 100, 000 cells per well in 12-well plates (Corning^®^ CellBIND^®^) and treated as abovementioned. After HG and HI conditions, cells were washed with PBS to remove dead cells and debris. Cells were fixed, permeabilized, and blocked as described before [17]. Next, cells were incubated with a polyclonal antibody against the vWf (1:400, 3% BSA in PBS) overnight at 4°C and thereafter washed 3 x with PBS. Alexa Fluor 488-labeled anti-rabbit (1:400 in PBS) was then used as a secondary antibody for 1 h at RT and washed 3 x with PBS. As a negative control, cells were incubated only with secondary antibody to assess for non-specific binding. Cell nuclei were stained with Hoechst 33258 (2 µg/ml in HBSS) as described above. Finally, fluorescence images were taken in at least three random fields per condition using an EVOS^®^ FLoid^®^ Cell Imaging Station with a fixed 20x air objective. Image analysis was performed by ImageJ software (version 2.0.0).

### 2.5 Assessment of mitochondrial inner mass and membrane potential

Cells were seeded at 100,000 cells per well in 12-well plates (Corning^®^ CellBIND^®^) and treated as above. After HG and HI conditions, cells were washed with PBS to remove dead cells and debris. Cells were washed with PBS and incubated with either 200 nmol/L MitoTracker Green FM or MitoTracker Red FM (in HBSS) for 30 min at 37 °C. Cells were washed with PBS and fluorescence images were immediately taken using an EVOS^®^ FLoid^®^ Cell Imaging Station with a fixed 20x air objective. The quantification of the mitochondrial mass and membrane potential was done by calculating the corrected total cell fluorescence (CTCF) tool from Image J software (version 2.0.0) using the following equation: Integrated density-(Area of selected cell x mean fluorescence of background readings) [17]. Two random fields per condition were chosen to capture images.

### 2.6 Sample preparation for MS analysis

Cells were seeded at 300,000 cells per well in 6-well plates (Corning^®^ CellBIND^®^) and treated as above. After HG and HI conditions, cells were washed 3x with PBS to remove dead cells and debris. Proteins were extracted, reduced, alkylated and digested using the Pierce™ Mass Spec Sample Prep Kit for Cultured Cells (Cat. No. 84840) as per the manufacturer’s instructions. Tryptic peptides were dried down by SpeedVac and reconstituted in water/acetonitrile 95:5 v/v with 0.1% formic acid. The peptide concentration was analyzed by using the Pierce™ Quantitative Colorimetric Peptide Assay kit (Cat. No. 23275) as per the manufacturer’s instructions.

### 2.7 MS analysis

Peptides were analyzed by reverse-phase high-pressure liquid chromatography (LC) coupled with electrospray ionization (ESI) tandem MS (LC-ESI-MS/MS). LC was carried out using an Eksigent nanoLC^®^ 400 system (AB Sciex, Foster City, CA, USA) with a HALO Fused-Core C18 (0.3 × 150 mm, 2.7 μm, 90 Å pore size, Eksigent AB Sciex, Foster City, CA, USA). The mobile phases A=0.1% formic acid in water and B=0.1% formic acid in acetonitrile were used. The elution of peptides was achieved by using a multistep linear gradient starting at 5% solvent B for 5 min then increased to 10% solvent B during 35 min, followed by an increase to 30% solvent B during 60 min, and a steep increase to 100% B for 35 min and held constant for 15 min. Six minutes post run with mobile phase A were applied to ensure column re-equilibration. The flow rate was 5 μL/min, and the sample injection was 4 μL. Two blanks were run between sample injections to minimize potential carryover. The eluate from the LC was delivered directly to the TurboV source of a TripleTOF 5600+ mass spectrometer (AB Sciex, Foster City, CA, USA) using ESI under positive ion mode. ESI source conditions were set as following: IonSpray Voltage Floating (ISVF), −5500 V; source temperature, 450 °C; Curtain gas (CUR), 30; Ion Source Gas 1 (GS1), 25; Ion Source Gas 2 (GS2), 35.

To create the SWATH-MS spectral library, the tryptic peptides from all samples (NG [n=3] and HG+HI [n=3]) (1 µg) were pooled and subjected to LC as abovementioned, and data-dependent acquisition (DDA) was employed. The mass spectrometer was operated in information-dependent acquisition with high sensitivity mode selected, automatically switching between full-scan MS and MS/MS. The accumulation time for TOF MS scan was 0.25 s/spectra over the m/z range 350-1200 Da and for MS/MS scan was 0.05 s/spectra over the m/z range 50-1200 Da (cycle time 1.3 s). The IDA settings were: charge state +2 to +4, intensity 150 cps, exclude isotopes within 6 Da, mass tolerance 50 mDa and a maximum number of candidate ions 20. Under IDA settings, the ‘‘exclude former target ions’’ was set as 20 s after two occurrences and ‘‘dynamic background subtract’’ was selected. The collision energies were varied to optimize sensitivity. The instrument was automatically calibrated by the batch mode using appropriate positive TOF MS and MS/MS calibration solutions before sample injections and after injection of two samples (<3.5 working hours) to ensure a mass accuracy of <5 ppm for both MS and MS/MS data.

For data-independent acquisition (DIA), fifty-six precursor isolation variable windows were generated using the SWATH Variable Window Calculator (AB Sciex, Foster City, CA, USA) based on precursor *m/z* frequencies in the DDA samples, with a minimum window width of 5 *m/z* (**Supporting Information Table 1**). The accumulation time for TOF MS scan was 0.100 s/spectra over the m/z range 350-1200 Da and MS/MS data were acquired from 350-1200 Da with an accumulation time of 0.05 s/spectra per SWATH variable window (cycle time 2.9 s). Other parameters were the same as those used in DDA.

### 2.8 MS data mining

Data from DDA samples were processed using ProteinPilot software version 4.2 (AB Sciex, Foster City, CA, USA) with the Paragon algorithm. MS/MS data were searched against a SwissProt Bos taurus database containing 6 006 reviewed proteins. The parameters used were as follows: Sample type, Identification; Cys Alkylation, Iodoacetamide; digestion, trypsin; Instrument, TripleTOF 5600; Special Factors, none; Species, Bos Taurus; ID Focus, Biological modifications, and amino acid substitutions; Search Effort, Thorough ID. False discovery rate analysis was also performed.

Targeted data extraction of SWATH files was done using the SWATH^®^ Acquisition MicroApp 2.0 in PeakView version 1.2 (AB Sciex, Foster City, CA, USA) along with the spectral library generated by DDA. Retention time calibration among samples was done manually using two endogenous peptides (>10,000 intensity) every 10 minutes throughout the LC gradient. The parameters used were as follows: Number of peptides per protein, 6; Number of transitions per peptide, 6; Peptide confidence threshold, 99%; False discovery rate threshold, 1%; exclude modified peptides, checked; XIC extraction window, 10 min, and; XIC width, 75 ppm. Protein peak areas were exported to Markerview (AB Sciex, Foster City, CA, USA) for further processing. Protein areas were normalized by the Total Area Sums algorithm in Markerview and exported to Microsoft Excel version 15.13.3 for statistical analysis. The reversed and common contaminants hits were removed before further processing. Proteins with a fold change ≥ 1.3 or ≤ 1/1.3 and a *p-value* <0.05 (Welch’s t-test) were considered differentially abundant between NG and HG+HI conditions. The mass spectrometry proteomics data have been deposited to the ProteomeXchange Consortium via the PRIDE [18] partner repository with the dataset identifier PXD013643.

### 2.9 Bioinformatic analysis

Gene ontology enrichment analysis of biological process and molecular function annotations were done using ClueGo app for Cytoscape (version 3.7.0) [19]. An enrichment/depletion (two-sided hypergeometric test) method with Benjamini-Hochberg correction was performed. A minimum and maximal GO level of 3 and 8 were used, respectively. Kappa Score was set to 0.4.

### 2.10 Statistical analysis

Data normality distribution was analyzed by the D’Angostino-Pearson normality test using PRISM 6. Select data were normalized to the respective control median values and are expressed as median (10^th^ percentile to 90^th^ percentile) (unless otherwise stated) derived from at least three independent experiments performed each in triplicate. Statistical analysis of data was performed by the Mann-Whitney test. A p<0.05 was considered statistically significant. Graphs were created and analyzed using PRIMS 6.0 (GraphPad Software, San Diego, CA).

## Results

An *in vitro* model of diabetic coronary endothelium was utilized by following the scheme illustrated in **Fig. 1**. First, insulin resistance was induced by treating the endothelial cells with a high concentration of insulin (HI, 100 nmol/L) for 3 days [15] followed by high-glucose (HG, 20 mmol/L) and HI for 9 days. A relatively long period with HG was chosen to avoid the acute effects of HG on cell proliferation (<48 h) [20]. Also, cells were maintained in a media with reduced serum to keep them under a quiescent mode. Endothelial cells with more than 2 nuclei were 3 times more abundant in HG+HI conditions compared to NG conditions and presented with variable size and shape (**Fig. 2A, 2B and 3A**). MVECs had a higher nucleic acid content (7.2-Fold, **Fig. 3D**), cell diameter (2.2-Fold, **Fig. 3B**), and cell area (11.4-Fold, **Fig. 3C**) than mononuclear cells. The cell diameter of MVECs and mononuclear cells ranged from 38 to 261 µm and 25 to 169.2 µm, respectively (**Supplementary Fig. 1**).

**Fig 1.**
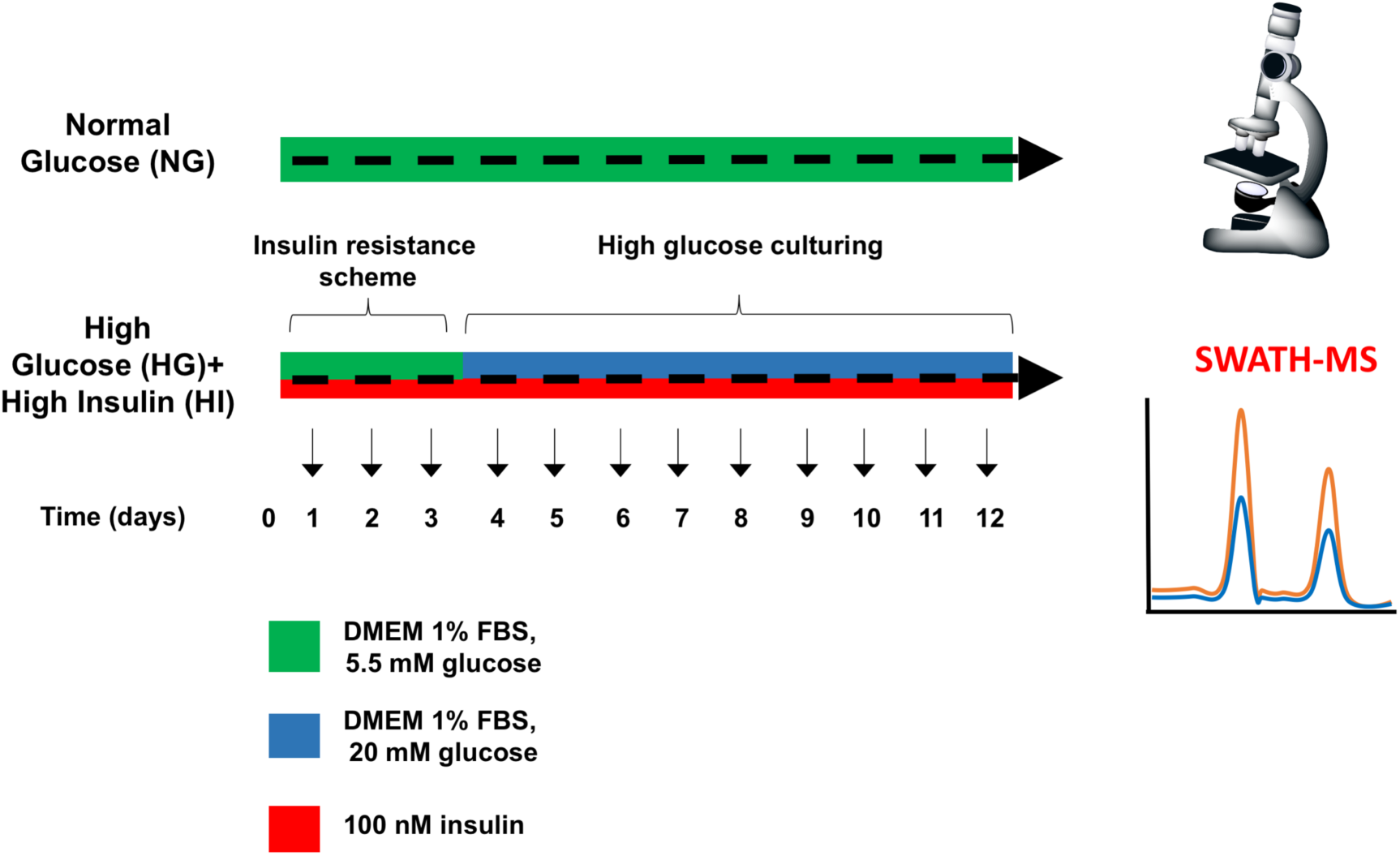
Illustration of the methodology followed in this study.

**Fig 2.**
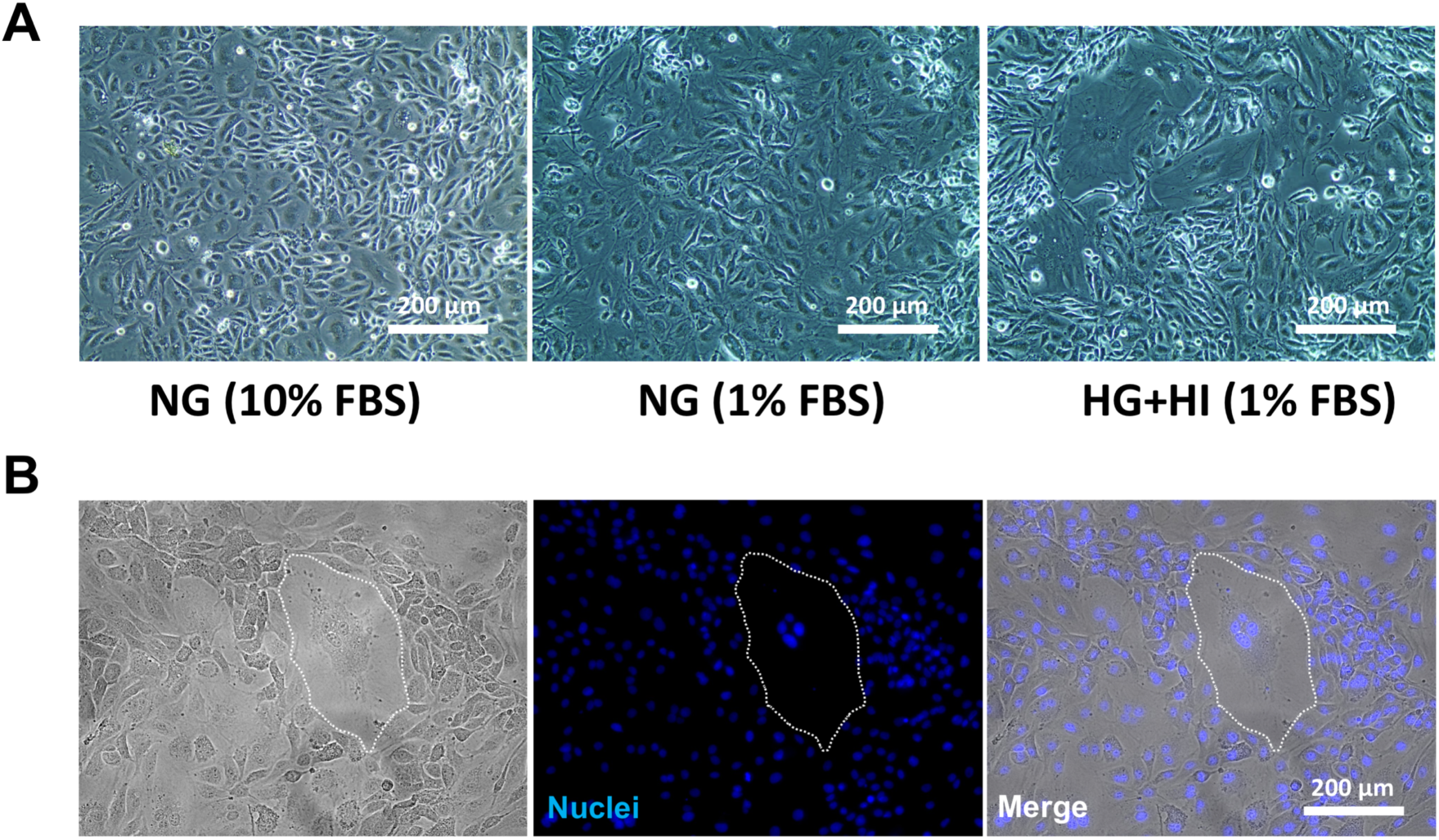
Presence and morphology of multinucleated variant endothelial cells (MVECs) in coronary artery endothelium cultured under simulated diabetes. (A) Representative transmitted-light micrographs of bovine coronary artery endothelium cells (BCAECs) cultured in either regular growth media (DMEM, 5.5 mmol/L glucose, and 10% FBS), reduced serum media (DMEM, 5.5 mmol/L glucose [NG], and 1% FBS) or diabetic media (DMEM, 20 mmol/L glucose [HG] plus 100 nmol/L insulin [HI], 1% FBS). Refer to the methods section for a more detailed description. (B) Representative image of a MVEC in diabetic media. Nuclei were stained with Hoechst 33258 for 30 minutes.

**Fig 3.**
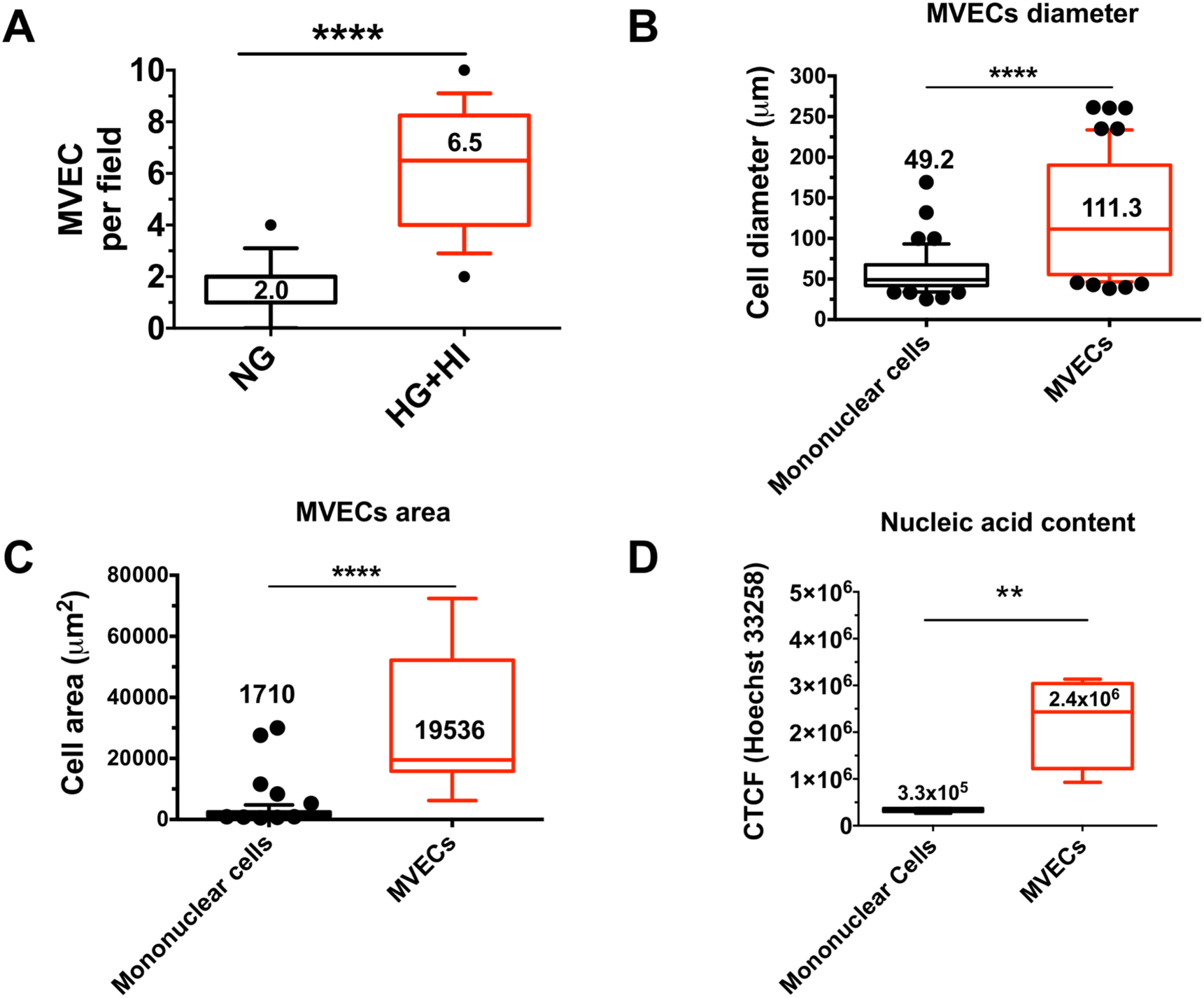
Abundance quantification and morphological characterization of variant endothelial cells (MVECs) in coronary artery endothelium cultured under simulated diabetes. Bovine coronary artery endothelial cells (BCAECs) were cultured in either normal glucose media (NG, 5.5 mmol/L) or high-glucose (HG, 20 mmol/L) plus high-insulin (HI, 100 nmol/L) media for nine days. Refer to the methods section for a more detailed description. (A) Abundance of MVECs in NG and HG+HI media. (B) Quantification of the diameter and (C) area of MVECs and mononuclear cells cultured under simulated diabetes using ImageJ software tools. (D) Nucleic acid content of MVECs and mononuclear cells cultured under simulated diabetes. Nucleic acid staining was done using Hoechst 33258. The corrected total cell fluorescence (CTCF) was determined using ImageJ software. Data are expressed as median (10^th^ percentile to 90^th^ percentile) derived from at least three independent experiments performed each in triplicate. Statistical analysis of data was performed by the Mann-Whitney test. **p<0.01, ****p<0.0001.

To demonstrate an endothelial phenotype in MVECs, we analyzed the presence of vWf, a well-known marker of endothelial cells [21]. The typical intracellular localization of vWf was observed in MVECs and mononuclear cells (**Fig. 4A**). No significant non-specific staining was noted (**Fig. 4B**).

**Fig 4.**
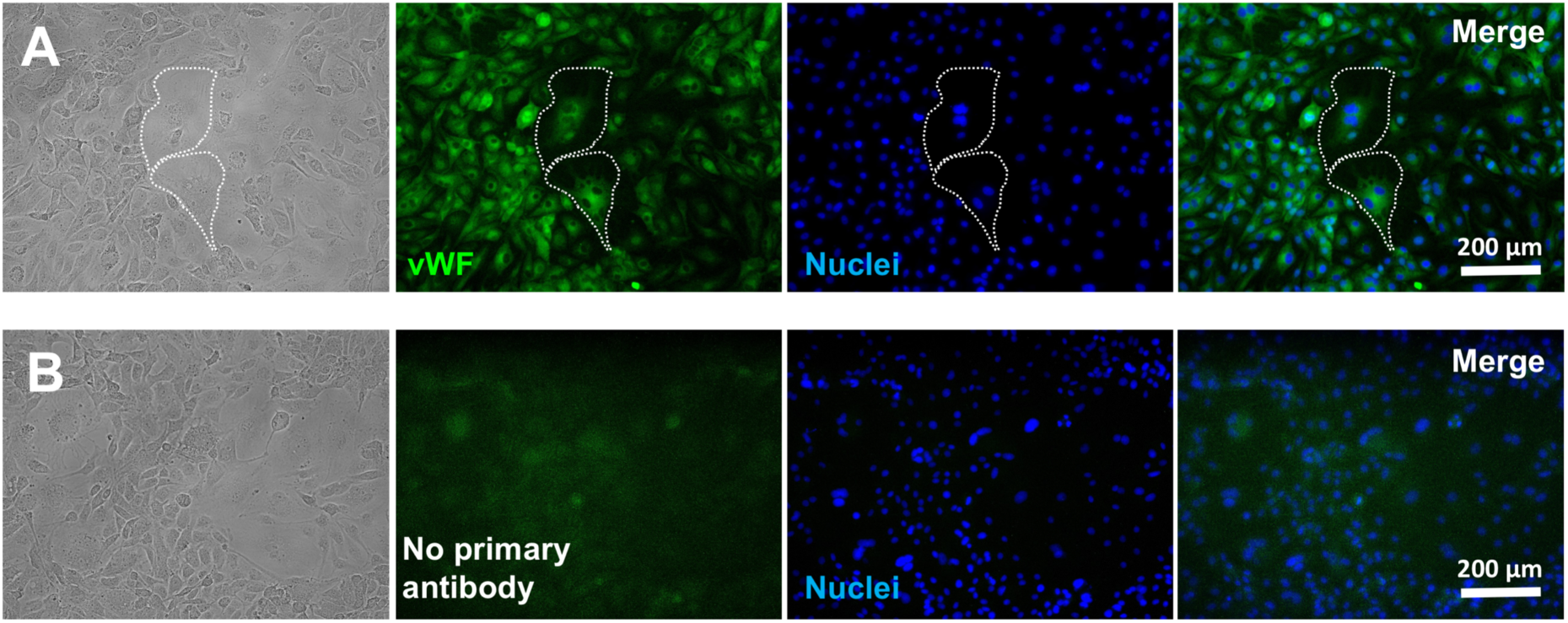
Presence of the von-Willebrand factor (vWf) in multinucleated variant endothelial cells (MVECs). (A) Representative immunofluorescent micrographs showing the localization of the vWf in fixed and permeabilized MVECs (green). Hoechst 33258 was used to stain the nuclei (blue). Intermittent white lines delineate the cell perimeter of a MVEC. (B) As a negative control, cells were only incubated with a secondary antibody to assess for non-specific binding. Refer to the methods section for a more detailed description.

Endothelial function is strongly influenced by mitochondria, and we and others have reported that HG triggers mitochondrial fragmentation or reduces mitochondrial biogenesis in cultured endothelial cells [22-24]. We therefore analyzed mitochondrial mass and membrane potential (a surrogate marker of mitochondrial function [25]) using mitochondrial selective fluorescent probes [26, 27] by fluorescence microscopy. As expected, MTG staining was significantly lower in mononuclear cells cultured in HG+HI conditions compared to those in NG conditions (**Fig. 5D and 5E**). However, MVECs showed a higher MTG staining compared to mononuclear cells (**Fig. 5A and 5B**). Contrary, MTR staining was higher in mononuclear cells cultured in HG+HI vs. NG conditions (**Fig. 5D and 5F**). MVECs showed a higher MTR staining compared to mononuclear cells similarly in magnitude to that observed with MTG (**Fig. 5A and 5C**). However, the ratio between MTR and MTG was 0.92 (**Supplementary Fig. 2**) suggesting that the increased MTR levels are attributed to the increase in mitochondrial mass rather than alterations at the mitochondrial membrane potential. Using quantitative MS-based proteomics, we next sought to determine the proteomic profile of endothelial cells grown under HG+HI vs. those grown in NG. A total of 939 proteins were quantified, wherein 14 and 25 proteins were significantly downregulated and upregulated in cells cultured with HG+HI vs. those in NG, respectively (**Fig. 6A**). To determine the biological context of the proteins differentially abundant between HG+HI and NG conditions, we performed a GO enrichment analysis filtering by molecular function, and biological process GO terms [19, 28]. Downregulated proteins were associated with the GO terms heparin binding and carboxylic binding (**Fig. 6B**). On the other hand, upregulated proteins were associated with the GO terms autophagy, positive regulation of intrinsic apoptotic signaling pathway, and platelet-activating factor acetyltransferase activity (**Fig. 6C**).

**Fig 5.**
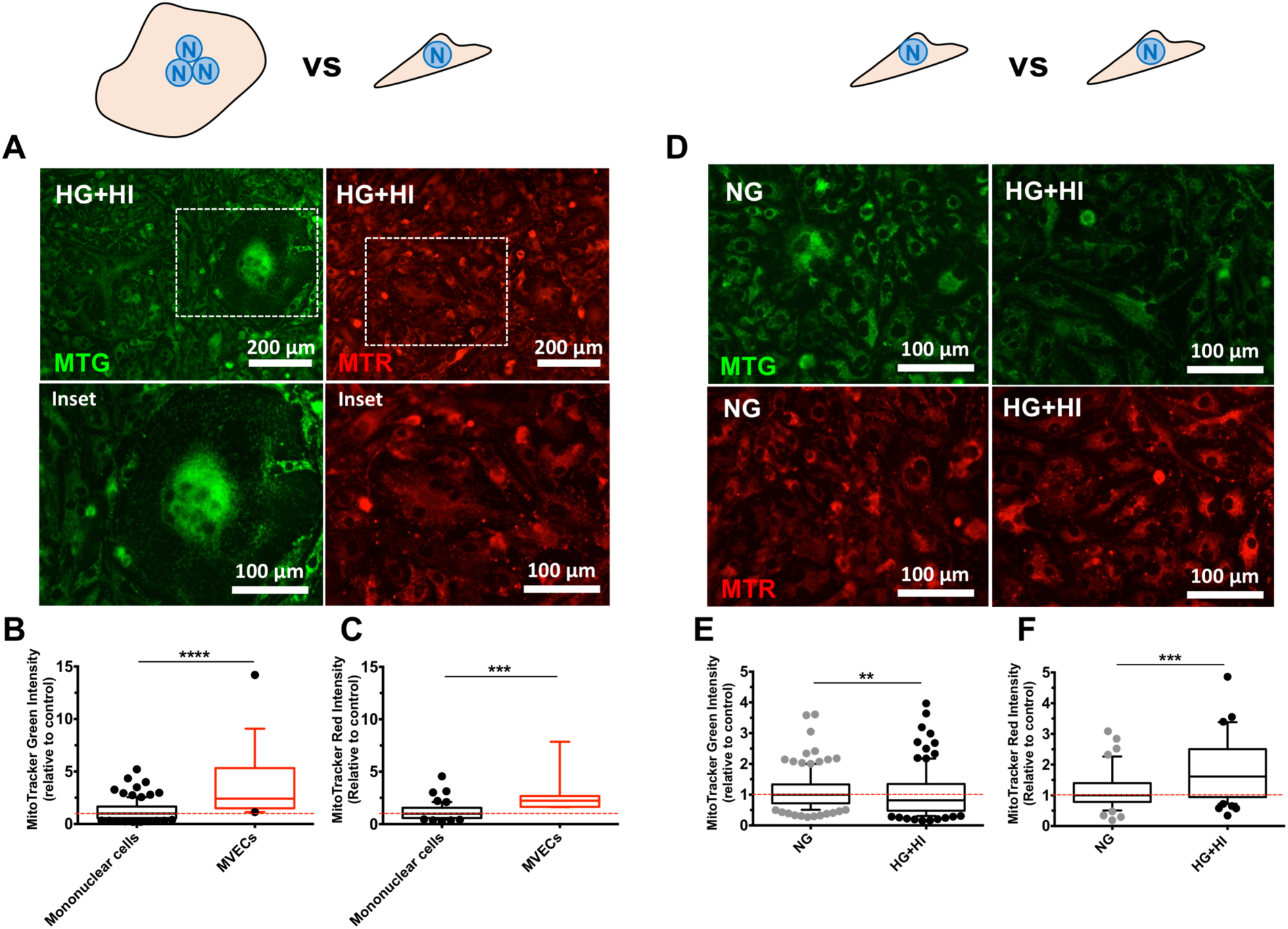
Mitochondrial structure and function of multinucleated variant endothelial cells (MVECs) and mononuclear cells cultured in simulates diabetes. Bovine coronary artery endothelial cells (BCECs) were stained with selective mitochondrial fluorescent probes to assess for mitochondrial mass (green) and mitochondrial membrane potential (red) levels. (A) Representative fluorescent micrographs of BCAECs cultured with high-glucose (HG, 20 mmol/L) plus high-insulin (HI, 100 nmol/L) media for nine days. White squares denote the presence of MVECs. Quantification of the mitochondrial mass (B) and mitochondrial membrane potential (C) of MVECs and mononuclear cells. (D) Representative fluorescent micrographs of BCAECs cultured with either normal glucose media (NG, 5.5 mmol/L) or high-glucose (HG, 20 mmol/L) plus high-insulin (HI, 100 nmol/L) media for nine days. Quantification of the mitochondrial mass (E) and mitochondrial membrane potential (F) of mononuclear cells cultured in NG and HG+HI media. Values in the group used as control were normalized and set as 1 (red intermittent line). Data are expressed as median (10^th^ percentile to 90^th^ percentile) derived from at least three independent experiments performed each in triplicate. Statistical analysis of data was performed by the Mann-Whitney test. **p<0.01, ***p<0.001, ****p<0.0001.

**Fig 6.**
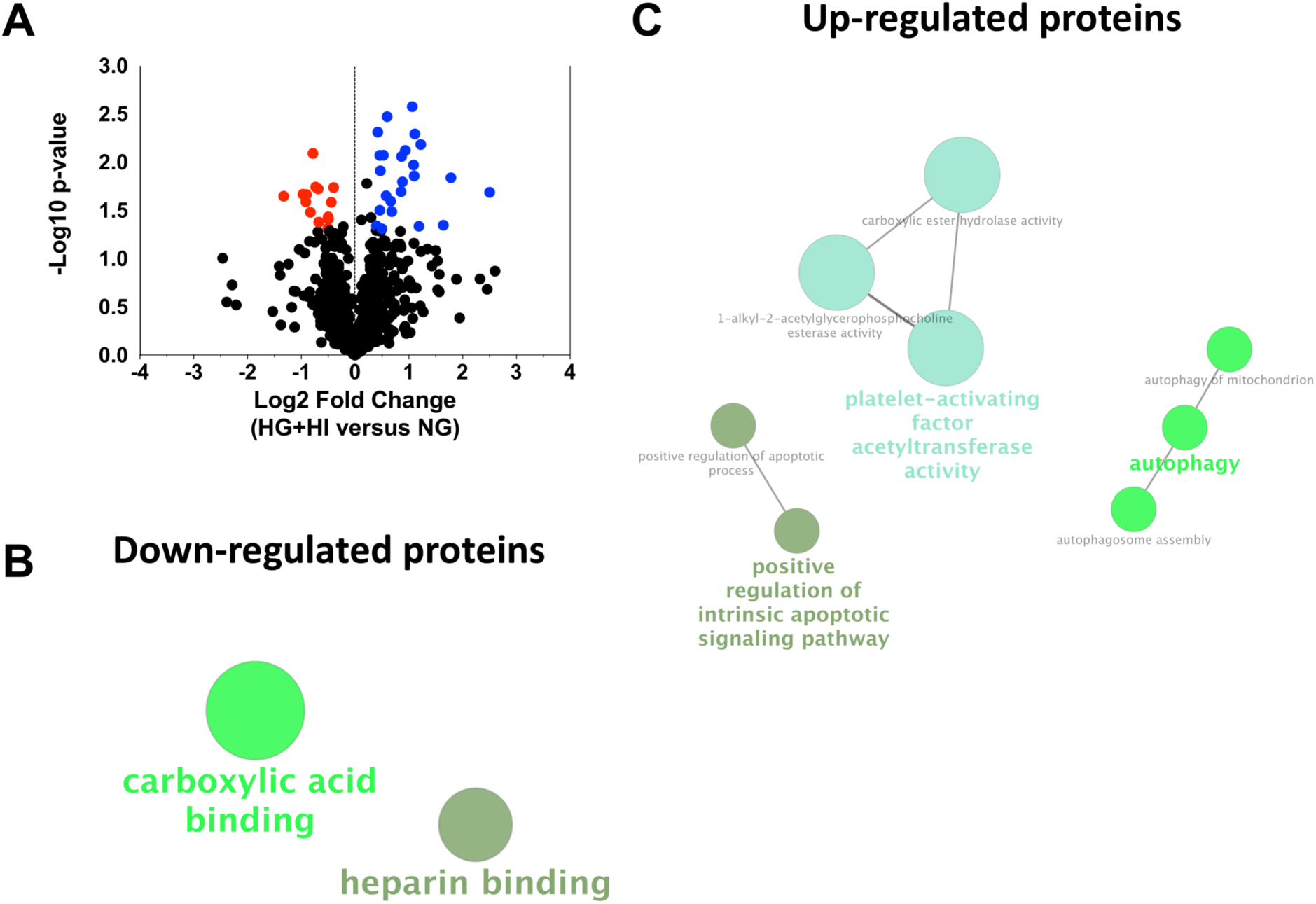
Quantitative proteomic analysis of bovine coronary artery endothelial cells (BCAECs) cultured in normal glucose (NG) and diabetic media (HG+HI). Volcano plot of all quantified proteins displaying differences in relative abundance between BCAECs cultured in NG (5.5 mmol/L glucose) media and diabetic media (HG+HI) for nine days. Values (dots) represent the HG+HI/NG ratio for all proteins. Red and blue dots denote downregulated and upregulated proteins in the HG+HI group vs. NG group, respectively. (B) Gene ontology (GO) enrichment analysis of downregulated proteins and (C) upregulated proteins, respectively. The name of the most significant term is shown in each group. The GO enrichment analyses were performed using biological process and molecular function annotations using ClueGo app for Cytoscape (version 3.7.0).

## 3. Discussion

Endothelial dysfunction in coronary endothelium is strongly linked to the development of myocardial infarction in subjects with T2DM [2]. Previous studies have demonstrated the role of HG in inducing mitochondrial damage in cultured endothelial cells [22-24], which is associated with endothelial dysfunction. To our knowledge, all published studies have focused on mononuclear endothelial cells, but there is evidence that a distinct endothelial phenotype exists in the vasculature [13]. These cells are multinucleated (>2 nuclei), bigger that mononuclear endothelial cells (referred to as MVECs), and present with altered functions (e.g., increased LDL uptake) [14]. The occurrence of MVECs in human aorta correlates with the severity of atherosclerosis [13] thereby suggesting a plausible role of these cells in the development of CVD. However, the presence or biological relevance of MVECs in the coronary endothelium remains obscure. Here, we have provided evidence of the existence of MVECs in cultured coronary endothelium, which is exacerbated by a simulated diabetic environment. The biological relevance of the multinucleation cellular process has been attributed to the stimulation of protective cellular mechanisms (e.g., immune cells) [29], senescence [30], and malignancy [31]. It is possible that MVECs occur in simulated diabetes as a compensatory mechanism to metabolize the excess of glucose or due to a damage in the mitotic machinery.

Moreover, simulated diabetes elicited differential effects on the endothelial mitochondria of mononuclear cells and MVECs. In agreement with various *in vitro* and *in vivo* studies, mononuclear endothelial cells showed reduced mitochondrial mass and increased mitochondrial membrane potential in HG environments [22-24, 32]. However, the opposite mitochondrial findings were noted in MVECs, which suggest differential mechanisms occurring among both endothelial phenotypes.

Our initial goal was to isolate MVECs by flow cytometry and cell sorting. However, we could not accomplish it due to several technical and biological limitations. Thus, to globally describe the cellular mechanisms altered by simulated diabetes, we performed a quantitative proteomic analysis on cells (mononuclear and MVECs mixed) cultured in NG and HG+HI and compared the protein profile amongst them.

Our proteomic analysis suggested the upregulation of proteins involved in mediating apoptosis and mitochondrial autophagy along with proteins associated with the suppression of inflammation. We particularly suggest critical roles of the platelet activating factor-acetylhydrolases (PAFAH1B2 and PAFAH1B3) [33] in mediating suppression of inflammation in endothelial cells under simulated diabetes. These findings suggest a balance between cellular damage and cellular protection mechanisms. Interestingly, we did not observe a considerable endothelial cell death even after nine days of simulated diabetes (data not shown), which suggests that endothelial cells may be resistant to the damage caused by a diabetic environment, at least after the period we tested [34]. On the other hand, we found a downregulation of molecular functions associated with angiogenesis and endothelial function in cells cultured with HG+HI. Specifically, we noted a decrease in the protein abundance of the connective tissue growth factor, a protein involved in the stimulation of angiogenesis [35], while in other tissues is upregulated by HG and associated with the development of fibrosis [36, 37].

In conclusion, we have evidenced that a diabetic environment leads to the formation of MVECs in coronary artery endothelium (**Fig. 7**). We also document on the differential effects that simulated diabetes has on the mitochondrial structure and function of mononuclear cells and MVECs. The unusual characteristics of these cells suggest that may play a role in the development and/or progression of vascular pathologies as those seen with T2DM.

**Fig 7.**
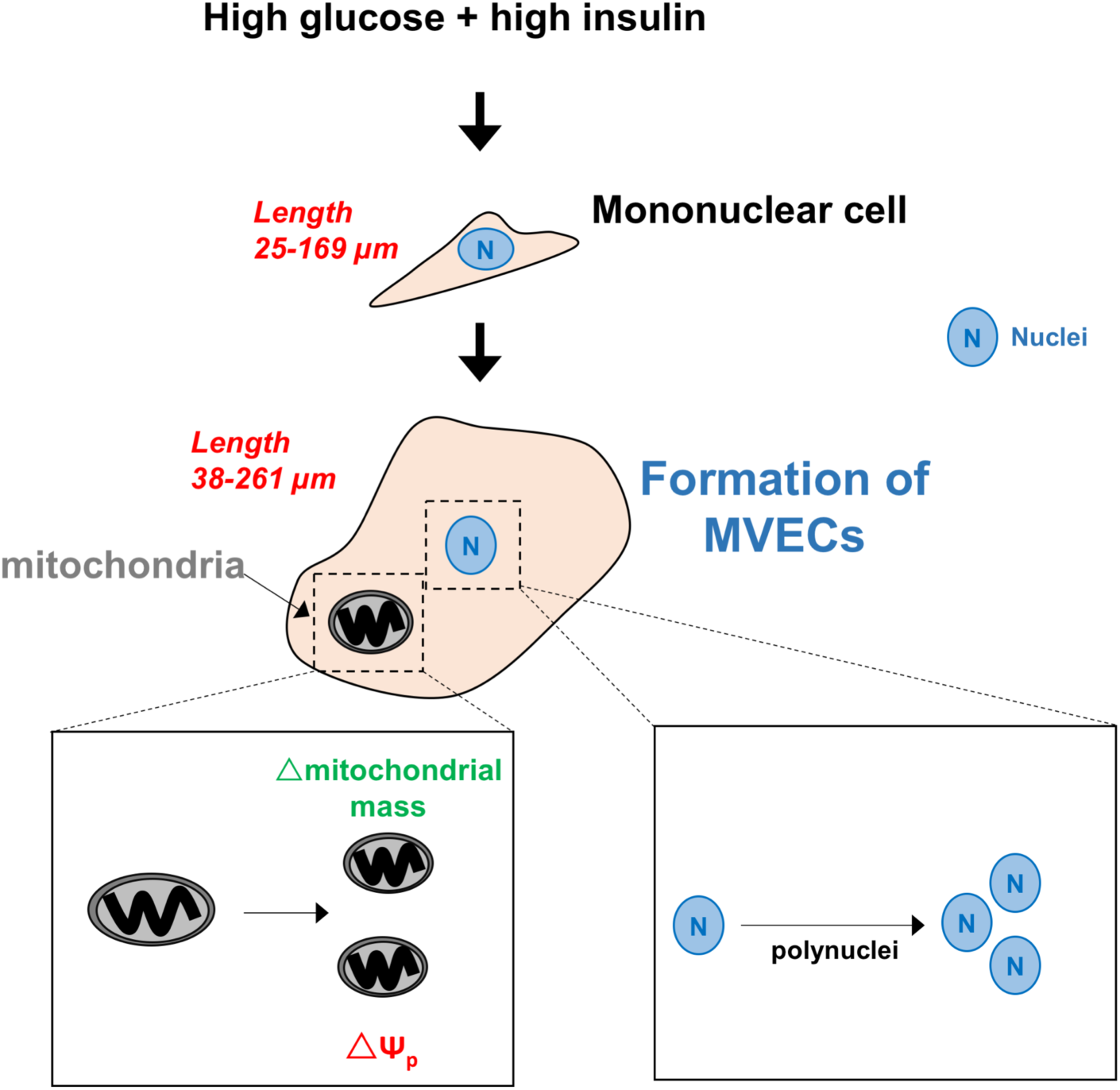
Summary illustration of study findings.

## 6. Acknowledgements

This work was derived from the Thesis Project of H.C.D.H. at the Posgrado en Ciencias de la Vida, CICESE.

