## Supplementary Information for "Formation of multinucleated variant endothelial cells with altered mitochondrial function in cultured coronary endothelium under simulated diabetes"

for

^3^ San Diego VA Healthcare System

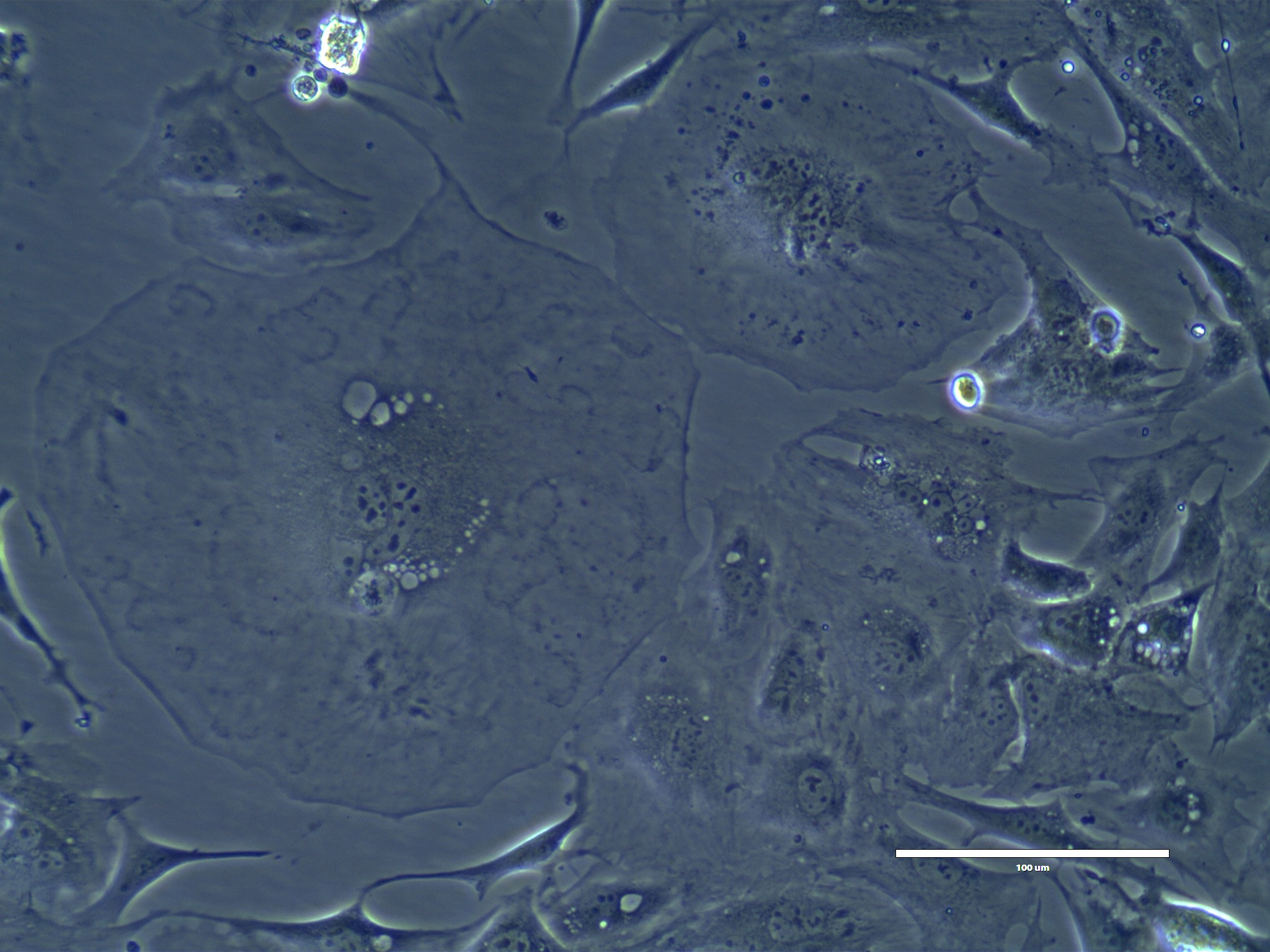

**Supplementary Fig. 1.** Representative transmitted-light micrograph of a multinucleated variant endothelial cell (MVEC).

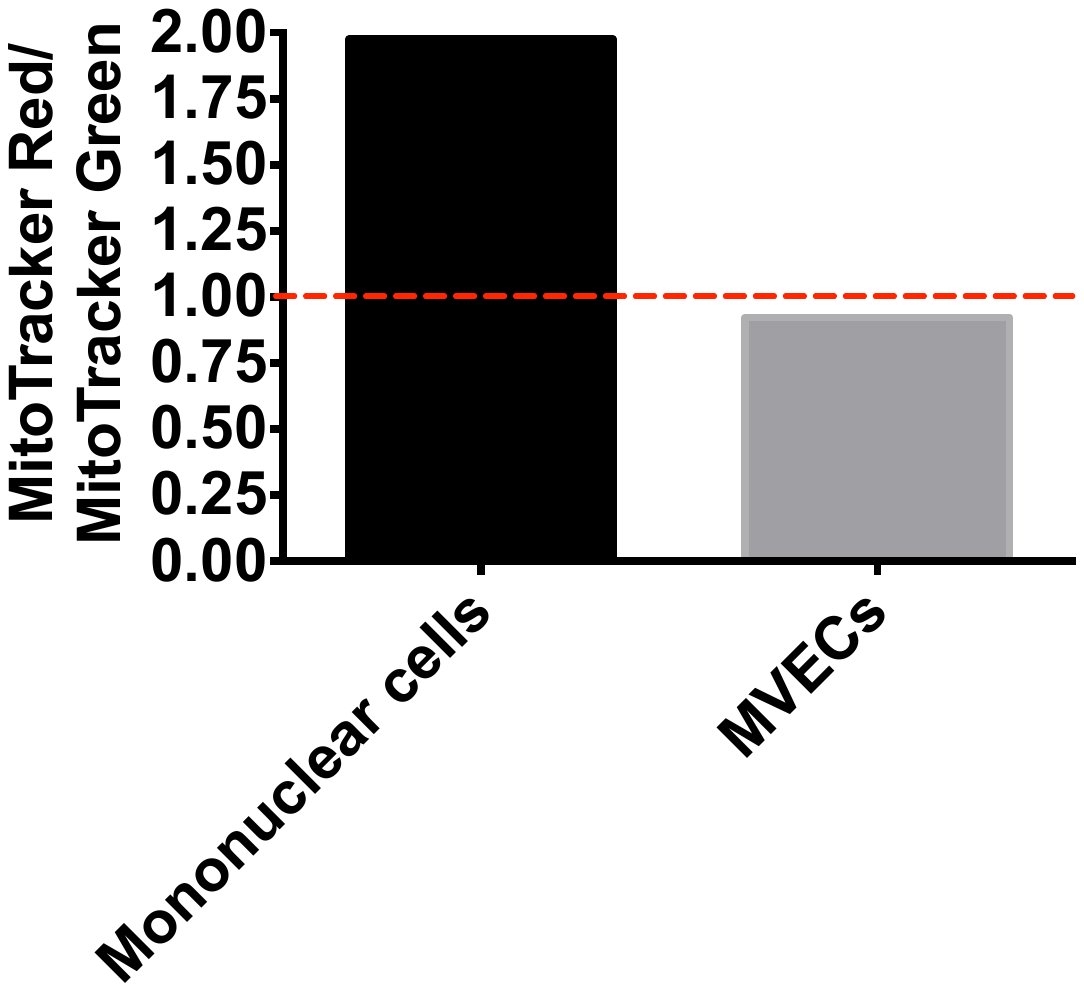

**Supplementary Fig. 2.** Ratio of the fluorescence levels between MitoTracker Red FM and MitoTracker Green FM of mononuclear cells and MVECs cultured in simulated diabetes.
